# Navigating the Maze: Identifying Potential Pitfalls in Attention State Classification from fMRI Brain Patterns

**DOI:** 10.1101/2024.12.20.629827

**Authors:** Ruben Andreas Bressler, Assunta Ciarlo, Sophie Raible, Giancarlo Valente, Michael Lührs, Ralph Tier, David E. Linden, Rainer Goebel

**Author notes:** shared first authorship (Ruben Andreas Bressler, Assunta Ciarlo).

## Abstract

Multi-voxel pattern analysis (MVPA) is a powerful technique to decode brain states from functional magnetic resonance imaging (fMRI) activity patterns. In neurofeedback (NF) applications, it has been used to perform real-time classification of brain activity patterns, establishing a closed-loop system that provides immediate feedback to the participants, enabling them to learn to control a complex mental state. However, MVPA has many potential limitations when applied to fMRI datasets (especially in real-time analysis) arising from small effect sizes, small number of training samples, high dimensionality of the data and, more generally, design choices. All these factors might produce inaccurate classification results. In this work, we followed a previous NF paradigm for sustained attention training. Participants were presented with composite images superimposing faces and scenes. They were instructed to focus on one class (either face or scene) for an extended period. A logistic regression classifier was trained to determine whether participants were adequately focusing on the instructed category based on their fMRI data. We analysed the classification outputs of the no-feedback training runs using various classifier settings, including whole brain data and different masking approaches, combined with different methods for the computation of single-trial fMRI responses. Furthermore, a ventricle mask was used as a control condition for the classification task, and simulations were carried out to assess the influence of the class order on the classification performances. We found inflation of the decoding accuracy for several common design choices and confounders. In particular, motion artefacts and low frequency drifts coupled with the task timing might have artificially increased the accuracy scores. Furthermore, the simulations revealed that fixed order in the presentation of experimental conditions resulted in further inflation of the classification accuracies especially in GLM-based and average-based trial estimate methods. We discuss the drawbacks of applying MVPA using the analysed sustained attention paradigm and provide insights for future improvements.

## Introduction

The intricate nature of the brain’s functional networks demands advanced techniques for signal analysis. Particularly, multi-voxel pattern analysis (MVPA) has been widely used to search for reproducible spatial patterns of activity that exhibit consistent distinctions across experimental conditions (Haynes, 2015; Mahmoudi et al., 2012; Weaverdyck et al., 2020). In fMRI, MVPA is usually implemented with machine learning (ML) models, which examine the covariance structure across voxels, rather than changes in the activation level of each voxel independently (Mur et al., 2009). Furthermore, it is sensitive to alterations in spatial activity patterns within individual brain regions as well as across entire brain networks. In the ML framework, MVPA reduces to a classification problem where a classifier is employed to capture the relationship between fMRI spatial activity patterns and experimental conditions (J. D. Cohen et al., 2017; Mahmoudi et al., 2012; Weaverdyck et al., 2020). Using ML terminology, fMRI patterns are *samples* on which the classification task is performed and usually consist of activation maps for each experimental trial (*single-trial estimates*). Brain voxels are *features* whose values define the activation level at a certain location, whereas the experimental conditions represent different *classes* to discriminate.

Linear classifiers, such as support vector machine (SVM), linear discriminant analysis (LDA), and logistic regression (LR), are the most widespread tools for fMRI pattern analysis (Mahmoudi et al., 2012). Additionally, non-linear classifiers, such as deep neural networks (DNNs), have gained attention as a means to solve complex classification problems (Lecun et al., 2015; Liang et al., 2023). However, DNN models tend to have limited interpretability, making it challenging to pre-determine the most suitable DNN architecture for a specific task (Guo et al., 2016) and require very large sample sizes to obtain good performance (Smucny et al., 2022). All these models have been used for the classification of functional brain data (De Martino et al., 2008; DeBettencourt et al., 2015; Mandelkow et al., 2016; X. Wang et al., 2020) and were usually trained in a supervised manner. For linear classifiers, supervised training consists of computing feature weights that correspond to the relative contribution of the features to the classification task, using a subset of the data (training set) with known class labels. After tuning the feature’s weight on the training set, the ML model is presented with new data (test set) and determines the most likely category label thereof, based on the prior training. The goodness of the prediction is frequently expressed as the proportion of correctly classified samples within the test set, often referred to as *the accuracy* score.

MVPA has also been used to decode brain signals in real-time. While the first development was based on a finger tapping task (LaConte et al., 2007), MVPA has been expanded further in the field of brain-computer interface (BCI) and neurofeedback (NF) (Sorger & Goebel, 2020). In BCI applications, participants learn to directly control an external device (e.g., communication, robotic arm) with their brain states. This concept can be extended to NF, wherein participants receive feedback from a measure of their brain activity and eventually learn to enhance their self-regulation strategies. For NF applications, MVPA approaches have been used to identify mind-wandering and focused states (McDonald et al., 2017), decoding of unconscious fear responses and treatment thereof (Taschereau-Dumouchel et al., 2018), classification of emotional states (Sitaram et al., 2011), and controlling an avatar (O. Cohen et al., 2014).

However, to reliably use MVPA on fMRI data, it is important to be aware of certain limitations. First of all, the combination of high spatial resolution, which results in a high number of voxels/features, and limited availability of experimental trials in fMRI datasets makes the ML models prone to overfitting (Jamalabadi et al., 2016; Smith et al., 2011), especially when applied across the whole brain. Therefore, it has been advised to reduce the dimensionality of the data, which could also help to exclude non-informative features that might degrade MVPA performances (Craddock et al., 2012; De Martino et al., 2008; Madsen et al., 2017). When a priori information is available, one possible approach for feature reduction is masking. With masking, only certain voxels are included as features for the MVPA based on their anatomical location and/or associated cognitive functions. Functional masks can also be obtained from independent experiments specifically designed to localise brain regions responsive to a task of interest (Jamalabadi et al., 2016). Additionally, alternative univariate and multivariate approaches for feature selection have been developed (for a more comprehensive description see Formisano et al., 2008; L. Wang et al., 2013). The dual problem that arises from a small sample size, due to the limited number of participants and/or a small number of experimental trials available for each participant, leads to inaccurate characterisation of the class properties. Furthermore, class differences might be hard to detect because of the low signal-to-noise ratio or small effect size of the fMRI responses for each experimental trial. To address these issues, different methods for the computation of single-trial responses have been devised. In literature, many studies used patterns derived from voxel-level beta estimates or corresponding t-values of a general linear model (GLM) as input for the classification task (De Martino et al., 2008; Madsen et al., 2018). To maximise the number of trials, fast event-related designs are usually employed with rather short inter-trial intervals, which results in temporal correlations between estimated activity patterns of neighbouring trials (Mumford et al., 2014). For those cases, other GLM-based approaches have been developed specifically to better denoise/disentangle single-trial estimates (for further details see Abdulrahman & Henson, 2016; Prince et al., 2022). The advantage of GLM-based approaches for the estimation of single-trial responses is that confounds, such as drifts and motion parameters, can be added to the model, resulting in a more reliable pattern computation. Several studies also used raw fMRI data as input of the ML model (typically after percent signal change transformation or z-scoring), defining the patterns by single time points or temporal block averages (Baucom et al., 2012; Cox & Savoy, 2003; DeBettencourt et al., 2015). It is important to note that different methods for estimating response patterns could yield differences in decoding performance, and that, therefore, an accurate method selection, tailored to the specific study design, is needed (Misaki et al., 2010).

In addition to the different trial response estimation techniques and feature selection methods, the classification accuracy is also influenced by other factors associated with the overall study design. Certain design choices, such as the absence of randomization in the order of the experimental conditions, might introduce biases into the data that the classifier could potentially detect, rather than, or on top of, the actual class differences. More generally, any structure in the presentation order of the experimental conditions could result in an inflation of the observed difference between classes and even lead to above chance classification accuracy in the absence of any real effect (Hebart & Baker, 2018; Mumford et al., 2014).

Furthermore, the presence of noise due to motion or physiological drifts could also alter the real pattern structure, leading to additional unintended biases that the classifier might detect. Although it is standard practice to use nuisance regressors for denoising BOLD data in preparation for functional connectivity or univariate analysis, the impact of motion or physiological fluctuations on multivariate decoders is often disregarded, and only a few studies have investigated this effect and/or proposed denoising solutions (Prince et al., 2022; Stehr et al., 2023). Taken together, all these factors could bias the classifier output and eventually lead to erroneous results. Consequently, a meticulous examination of these factors is imperative, particularly in the case of real-time fMRI applications where, due to time constraints, the number of available trials is even more limited, along with the possibilities for data denoising.

In this work, we applied MVPA to a dataset acquired by adapting the design of a previous real-time fMRI NF study (DeBettencourt et al., 2015) and investigated the influence of multiple factors on the classifier performance. In the previous study, MVPA-based NF was successfully applied to train sustained attention. Indeed, MVPA is particularly well suited for investigating cognitive processes that involve distributed brain networks, such as sustained attention (Fox et al., 2005, 2006; Power et al., 2011). The sustained attention network (SAN) includes the intraparietal sulcus (IPS), the inferior frontal junction (IFJ) and the anterior midcingulate cortex (aMCC). Within the attention network, functioning is heterogeneously distributed, i.e., there is no central hub in control of all other areas (Langner & Eickhoff, 2013; Posner & Dehaene, 1994). It has been proposed that, with the application of MVPA, it is possible to decode attentional states from functional brain imaging signals related to sustained attention. DeBettencourt et al. performed real-time decoding of attentional states to provide immediate feedback to the participants during NF training. In the original study design, participants were presented with composite images superimposing faces and scenes and were instructed to focus on one class (either face or scene) for a prolonged period.

By using regularised LR based on whole-brain data, it was determined whether participants were adequately focusing on the instructed class at a given time. The proposed paradigm included long task periods of 50 seconds to effectively train sustained attention, which might be particularly vulnerable to low frequency fluctuations and inevitably leads to a small number of training samples when block approaches are used for the computation of single-trial responses. We employed a similar experimental paradigm, focusing on the analysis of the classifier training runs, and evaluated the impact of different trial response estimation methods and different masks, relevant for the sustained attention task, on the performances of a LR classifier. The influence of the fusiform face area (FFA) and parahippocampal place area (PPA) information on the classification accuracy was of particular interest as these areas are perceptually specialised for the target stimuli of the attention task, and both areas have been shown to be modulated by attention (Al-Aidroos et al., 2012; Baldauf & Desimone, 2014; Chee et al., 2010; Chiu et al., 2011; Downing et al., 2001; O’Craven et al., 1999; Reddy et al., 2007; Wojciulik et al., 1998; Yi et al., 2006). Classification from ventricle mask data was also performed as a control condition to detect potential non-task-related effects. Additionally, fMRI data simulation was carried out to specifically assess the influence of the order of the conditions on the classification accuracy. We aim to identify the influence of the masking approaches, different single-trial estimation methods, order effects, and confounding factors on the classifier’s classification accuracy.

## Methods

### Participants

The Ethics Review Committee Psychology and Neuroscience of Maastricht University reviewed and approved the study. 13 male recruits (mean age 34 years, age std = 2.6) of the Dutch police special forces (Dienst Speciale Interventies, DSI) participated in the study. All participants have previously served in the Dutch military (mean time served in the military ten years). All participants provided written informed consent before participating in the study. They were also informed that their training at the Special Forces was not influenced by their participation in the study.

### Design

The participants underwent six weekly neurofeedback sessions, except for six participants for whom the last two sessions were only five days apart. An anatomical scan was collected at the beginning of each session, followed by the functional acquisitions which included multiple runs for emotion regulation neurofeedback (not analysed in this work, reported in Bressler et al., 2023) and runs for sustained attention neurofeedback. In the first session, a functional localizer was conducted (Attention network localizer), based on which a subject-specific mask of the attention network was created. During each session, one classifier training run for the sustained attention neurofeedback was collected, after which the sustained attention neurofeedback was performed. In this work, only data from the classifier training runs were analysed, thus resulting in 6 training runs per participant.

### MRI

#### MRI acquisition

MRI images were collected using a 3T MAGNETOM MRI system (Prisma Fit, Siemens Healthcare, Erlangen, Germany), in combination with a 64-channel head coil. For anatomical scans, a magnetization-prepared rapid acquisition gradient echo (MPRAGE) sequence (TR=2300 ms, TE=2,89 ms, FoV=256 mm², voxel size=1 mm isotropic) was used. Functional images were collected with a BOLD-fMRI multi-band accelerated echo-planar imaging (EPI) sequence (volumes=496, TR=1 s, TE=31 ms, number of axial slices=48, voxel size=2.5 mm isotropic, matrix= 90*90, FoV=224 mm², multi-band factor=4, phase encoding direction=AP). Additionally, a shorter functional run of 5 volumes with opposite phase encoding direction was acquired right before the first functional run and used for correcting susceptibility-induced artefacts. Standard Siemens pulse oximeter and respiration belt were used to measure heart rate and breathing signals.

#### Attention network localizer

A letter n-back task with eight blocks of zero- and two-back conditions was used to construct an attention network mask. A task block lasted 20 seconds, during which 10 letters were presented centrally for 1s each, separated by 1s of fixation cross. Blocks were interspersed with 20s of rest during which the instructions for the next block were displayed. For the zero-back blocks, participants were instructed to click a button whenever they saw the letter “X”. In the two-back blocks, they were instructed to click whenever they had seen the same letter two steps prior (i.e., with one other letter in between). The letter “X” was excluded from the two-back condition.

#### Attention task (classifier training runs)

During the classifier training runs, transparently overlaid images of faces and scenes were presented. Image composition was stable at 50% face and 50% scene. At the beginning of each block, participants were instructed to focus on either the faces or the scenes present in the image. Each run consisted of four blocks per category, lasting 50s each. Blocks alternated between focusing on faces and scenes, and were separated by 7s of rest during which the instructions for the next block were displayed for 1s. A run started and ended with 20s of rest (Fig. 1).

**Figure 1.**
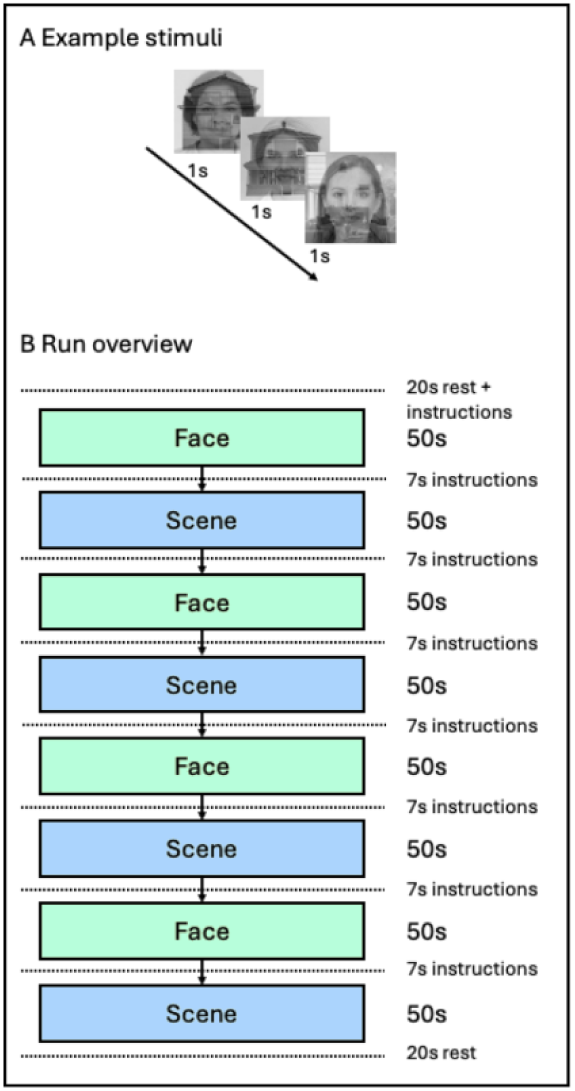
Example stimuli and classifier training run scheme. A shows examples of the composite images consisting of superimposed faces and scenes. The composition was stable at 50% face and 50% scene visibility. Stimuli were presented for one second. B shows a schematic run overview. Blocks of attending to faces and scenes were alternating. Each block lasted 50 seconds. Blocks were separated by seven seconds of rest during which the instructions for the next block were presented for 1 second. A run started and ended with 20 seconds of rest.

Faces could be either male or female, and scenes could be either indoors or outdoors. To control behaviourally for attention, an odd-ball task was designed. Participants were given instructions at the beginning of each block, informing them which of the four target categories (male/female for faces; indoors/outdoors for scenes) they had to respond to by button pressing. The respective face and scene target category was kept constant during a session. The target category for the odd-ball task appeared in 10% of all trials.

#### Data preprocessing

The functional data of the classifier training runs were preprocessed using BrainVoyager 22.2 (Brain Innovation, Maastricht, The Netherlands) and FSL 6.0.5.1 (http://www.fmrib.ox.ac.uk/fsl). The preprocessing steps included slice scan time correction, motion correction with sinc interpolation, using the first volume of each run as reference image, susceptibility distortion correction with the FSL *topup* routine, linear trend removal, and high-pass temporal filtering with a cut-off frequency of 0.008 Hz. To control for motion-induced artefacts, an outlier analysis was conducted over the spike count per run. A spike was defined as more than 0.25 mm root mean square (RMS) volume-to-volume displacement. All runs that fell outside the 75% percentile were excluded from the offline analysis. Additionally, functional runs with absolute maximum estimated motion greater than 5 mm/5° were also excluded.

The structural data of the first session of each subject were used as anatomical reference. Structural data preprocessing was performed with BrainVoyager and included inhomogeneity correction and skull stripping. Each preprocessed functional run was normalised to the Montreal National Institute (MNI) standard space by first aligning the functional data to the individual anatomical space, computing the transformation to the MNI space and resampling the functional data to 2mm isotropic resolution. Finally, the MNI-transformed functional data were spatially smoothed using a three-dimensional isotropic Gaussian kernel (FWHM=2mm).

### Analysis

#### Physiological Noise

Physiological noise data were visually inspected for data quality. Due to low data quality, 13% of the heart rate data and 50% of the breathing rate data of all runs had to be discarded (see supplementary S3). Due to the limited availability of usable physiological recordings, they were excluded from the subsequent analysis, as their impact on classification performance could not be reliably assessed.

#### Multivariate analysis

Multivariate analyses of the training runs were conducted at individual subject level, comparing different approaches for single-trial estimation and different masking methods. The classification task focused on the differentiation between brain activity patterns for the attention-to-face and attention-to-scene conditions computed for the attention blocks of each training run. The aim of this analysis was twofold: firstly, assess the impact of the choice of the trial estimation method on the classification performances; secondly, evaluate the influence of FFA and PPA spatial information.

The two-class classification was performed using a regularised logistic regression model (regularisation type = ‘L2’, penalty factor C = 1, solver= ‘liblinear’, max iteration number = 100) implemented in the *scikit-learn* python package (version 1.1.2) (Pedregosa et al., 2011). In section S2 of the Supplementary Materials, a comparison of the performances of the Python model with the Matlab implementation employed in the original paper (DeBettencourt et al., 2015) is reported. For both models, a leave-one-run-out cross-validation approach was used to assess the within-subject generalisation performances. Before training the classifier, the training data were (feature-wise) z-scored independently at each cross-validation step to avoid data leakage, and the computed transformation was applied to the test set of the same cross-validation level.

#### Single-trial estimation methods

Ten different methods for single-trial estimation were assessed. A detailed description of each method is provided in Table 1.

**Table 1.** Trial estimate methods details. GLM = general linear model; HRF = hemodynamic response function; BOLD = Blood Oxygen Level Dependent; TR = time repetition; PSC = percent signal change.

| Method | Description | Implementation Details |
| --- | --- | --- |
| **Single-trial t-values maps (*t-values*)** | Voxel-wise GLM contrast using a windowing approach. | Time window: 1s pre-onset, 55s post-onset of each attention block.<br>GLM regressors included a task-specific box-car predictor, convolved with a double-gamma HRF, alongside linear-trend and constant predictors. Z-scoring of data and predictors was performed before GLM fitting. 'Task vs. rest' contrasts were computed for each task block. |
| **Single-trial beta maps (*betas*)** | Voxel-wise windowed GLM focusing on effect size. | Similar approach to the *t-values* method, using the beta maps from GLM fitting. |
| **Single-trial t-values maps using motion confounds (MOCO *t-values*)** | Voxel-wise GLM contrast using a windowing approach and motion confounds. | Similar approach to the *t-values* method, including motion confounds in the GLM fitting. |
| **Single-trial beta maps using motion confounds (MOCO *betas*)** | Voxel-wise windowed GLM including motion confounds and focusing on effect size. | Similar approach to the *t-values* method, using the beta maps from GLM fitting including motion confounds. |
| **Single-trial average z-scored BOLD (*zscore-avg*)** | Average z-scored functional time series over a task block. | Functional time series were z-scored within each run. The average volume was computed over each task block, shifted by 4s to adjust for hemodynamic delay. |
| **Single-TR z-scored BOLD 10 samples**<br>(*zscore-singleTR-10*) | Uses single z-scored volumes as trials, selecting the last 10 volumes of each task block. | Functional time series were z-scored as in the *zscore-avg* method. The last 10 TRs of each trial block, after the 4s shift, were selected. |
| **Single-TR z-scored BOLD**<br>(*zscore-singleTR*) | Uses single z-scored volumes as trials. | Functional time series were z-scored as in the *zscore-avg* method. Single-trial estimates were obtained for each TR within a task block, adjusted for hemodynamic delay (4s shift). |
| **Single-trial average PSC BOLD**<br>(*PSC-avg*) | Average changes in BOLD signal from a global baseline over each task block. | Functional time series were PSC-transformed with respect to a global baseline, computed separately within each run by averaging the functional volumes of the first rest block and the time point preceding each task block. Single-trial estimates were computed averaging over each task block, adjusted for hemodynamic delay (4s shift). |
| **Single-TR PSC BOLD 10 samples**<br>(*PSC-singleTR-10*) | Uses single PSC-transformed volumes as trials, selecting the last 10 volumes of each task block. | Functional time series were PSC-transformed as in the *PSC-avg* method. The last 10 TRs of each trial block, after the 4s shift, were selected. |
| **Single-TR PSC BOLD**<br>(*PSC-singleTR*) | Uses single PSC volumes as trials. | Functional time series were PSC-transformed as in the *PSC-avg* method. Single-trial estimates were obtained for each TR within a task block, adjusted for hemodynamic delay (4s shift). |

#### Masking methods

Six different masks in MNI space were created for each subject and used to perform feature selection in the MVPA (Tab. 2).

**Table 2.**
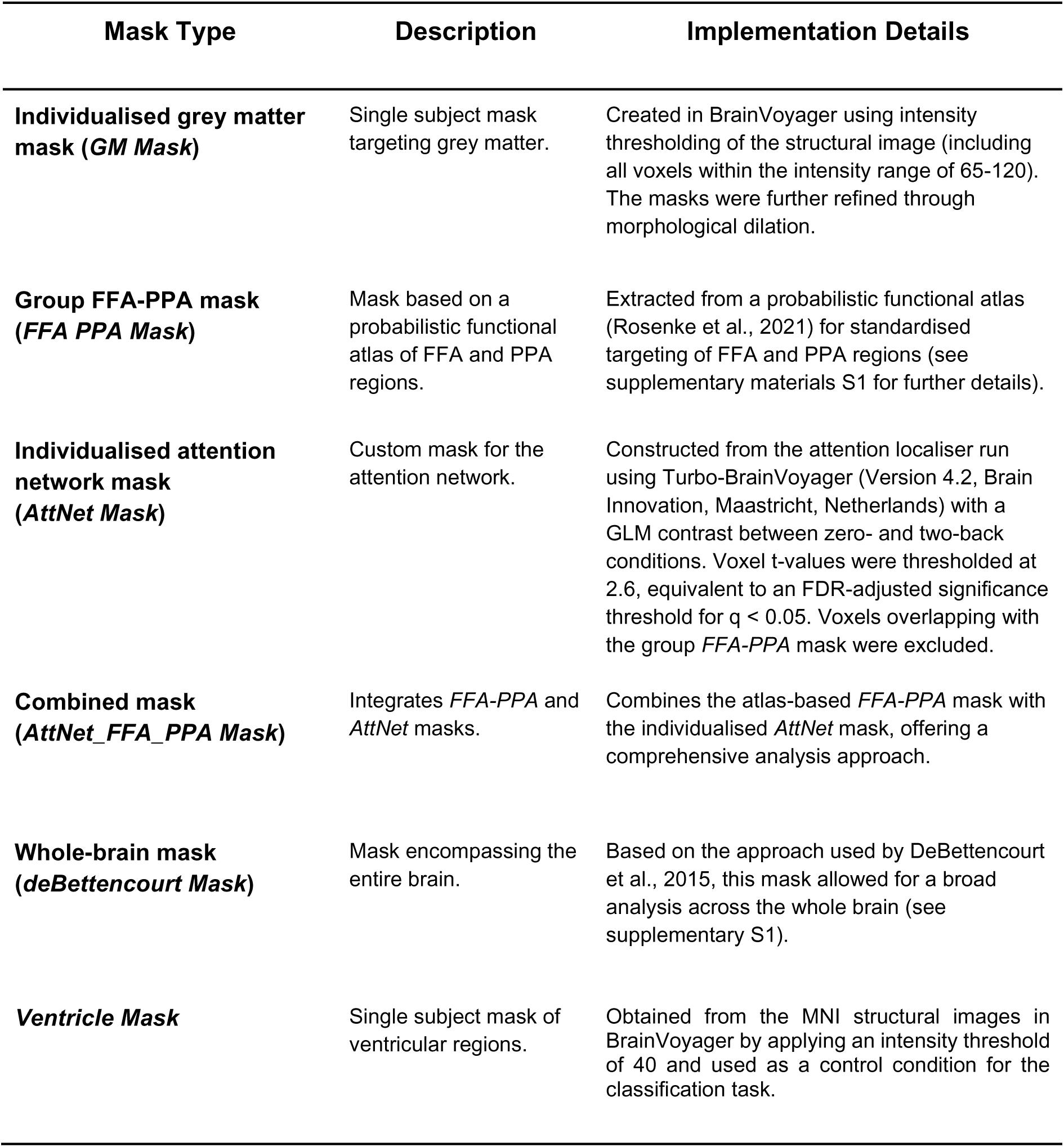
Masking methods details. FFA = fusiform face area; PPA = parahippocampal place area.

| Mask Type | Description | Implementation Details |
| --- | --- | --- |
| **Individualised grey matter mask (*GM Mask*)** | Single subject mask targeting grey matter. | Created in BrainVoyager using intensity thresholding of the structural image (including all voxels within the intensity range of 65-120). The masks were further refined through morphological dilation. |
| **Group FFA-PPA mask (*FFA PPA Mask*)** | Mask based on a probabilistic functional atlas of FFA and PPA regions. | Extracted from a probabilistic functional atlas (Rosenke et al., 2021) for standardised targeting of FFA and PPA regions (see supplementary materials S1 for further details). |
| **Individualised attention network mask (*AttNet Mask*)** | Custom mask for the attention network. | Constructed from the attention localiser run using Turbo-BrainVoyager (Version 4.2, Brain Innovation, Maastricht, Netherlands) with a GLM contrast between zero- and two-back conditions. Voxel t-values were thresholded at 2.6, equivalent to an FDR-adjusted significance threshold for $q < 0.05$. Voxels overlapping with the group *FFA-PPA* mask were excluded. |
| **Combined mask (*AttNet_FFA_PPA Mask*)** | Integrates *FFA-PPA* and *AttNet* masks. | Combines the atlas-based *FFA-PPA* mask with the individualised *AttNet* mask, offering a comprehensive analysis approach. |
| **Whole-brain mask (*deBettencourt Mask*)** | Mask encompassing the entire brain. | Based on the approach used by DeBettencourt et al., 2015, this mask allowed for a broad analysis across the whole brain (see supplementary S1). |
| **Ventricle Mask** | Single subject mask of ventricular regions. | Obtained from the MNI structural images in BrainVoyager by applying an intensity threshold of 40 and used as a control condition for the classification task. |

All masks except for the *ventricle* masks were further refined by removing noisy voxels. For each subject, voxels with mean intensity of the functional images lower than 100, calculated over all training runs, were excluded from the final masks. Additionally, all voxels outside an MNI brain mask obtained from the BrainVoyager MNI template were removed (except for the *deBettencourt* masks, see supplementary materials S1).

### Simulations

The original study design included a fixed order in the condition presentation for all training runs (i.e., the attention-to-face condition always preceded the attention-to-scene condition) that could potentially introduce problems for the decoding. To assess the effect of the condition order on the classification performance, time-series data were simulated using the same fixed-order design of the real data and a random-order design. No between-condition differences in the response amplitude were simulated to evaluate the effect of the design on the classification performance when no differentiation between the two conditions was expected. In the random-order design, the number of trials for each condition was kept constant within each run while the order of conditions was randomised independently for each run. BOLD time-series were simulated for each voxel within the group *FFA PPA mask* and the baseline value of each voxel was set to the average BOLD value computed across the first baseline block of all training runs and subjects. To simulate changes in the baseline level across different runs, random baseline shifts were added separately at each voxel. Percentage shift values were obtained from a zero-mean Gaussian distribution with a standard deviation of 10% for the FFA voxels and 5% for the PPA voxels, as estimated from the real data.

Ideal BOLD responses for the two conditions were obtained by convolving two box-car predictors with a standard double-gamma hemodynamic function (Friston et al., 1998) provided by the *‘spm’* model from *nilearn* python package (version 0.10.1) (Abraham et al., 2014; Pedregosa et al., 2011). For both the fixed- and random-order design, the amplitude of the ideal BOLD response was randomly drawn from a normal distribution ∼ *N(1, 0.1)* for all the task blocks, independently of the condition. Temporal fluctuations of the BOLD signal were obtained by adding auto-correlated Gaussian noise to the ideal time course at each voxel using the following autoregressive model (De Martino et al., 2008):

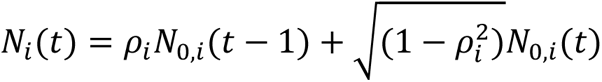

where 𝑁_0,𝑖_ is random Gaussian noise ∼ *N(0, 1),* 𝜌_𝑖_ ∼ 𝑁(0.5,1) is the autocorrelation coefficient at voxel *i*, and the scaling factor 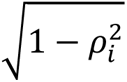 was applied to the base Gaussian noise so that the final noise signal had unitary variance. For simplicity, no covariance structure of noise was imposed across voxels. At each voxel, the ideal signal and the autocorrelated noise were combined after controlling for the task-baseline contrast-to-noise ratio (CNR). The CNR was defined as the ratio between the temporal standard deviation of the ideal signal and the temporal standard deviation of the noise (Welvaert & Rosseel, 2013) as indicated in the following formula:

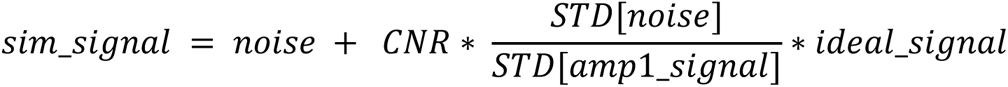

where 𝑎𝑚𝑝1_𝑠𝑖𝑔𝑛𝑎𝑙 is an ideal time course with fixed unitary maximum amplitude for each condition and was used to compute the signal scaling factor, whereas the 𝑖𝑑𝑒𝑎𝑙_𝑠𝑖𝑔𝑛𝑎𝑙 included random amplitude modulation. 1000 independent sets of autocorrelated noisy time series were generated, each one including 6 separate runs. The simulated data were generated for both fixed- and random-order design, with CNR values of 0.5, 1, 2 and 5 (an example is reported in section S5 of the supplementary materials). This procedure resulted in a total of 8000 simulated data sets, consisting of 1000 simulations per combination of design type and CNR level. The same autocorrelated noise was used while changing the design type and CNR level. Additionally, the amplitude modulation of the ideal signal was kept constant while varying the CNR.

A binary classification was performed using the same python logistic regression classifier and cross-validation procedure employed in the analysis of the real data.

Two-tailed one-sample t-tests were used to test against the null hypothesis that the obtained accuracy scores came from a normal distribution with a mean equal to 0.5 for both fixed- and random-order design, each trial estimate method and CNR value. The results were corrected for multiple comparisons using the Bonferroni criterion (n=64, where n is the product of multiplying the number of design types (2), by the number of trial estimate methods (8) and CNR levels (4)). Additionally, changes in the variance of the accuracy distribution across CNR values were also tested using Levene’s test (Levene, 1960) for each trial estimate method and simulated design, and results were corrected using the Bonferroni criterion (n=16). Statistical analyses were performed using Matlab 2021a (https://it.mathworks.com/products/matlab.html).

## Results

### Data exclusion

Due to excessive motion in most of the runs, one subject was completely excluded from the offline analysis. Furthermore, one run of participant P05 was excluded due to maximum absolute motion higher than 5 mm/degree, and one run of participant P06 was excluded due to partial coverage of the FFA and PPA regions.

### Multivariate analysis

The behavioural results of the oddball task showed that the participants were focused on the given task, exhibiting a mean response rate across all runs of 89.9%.

Considering the multivariate analysis of the training runs, figure 2 shows the average cross-validated accuracy scores across all subjects, obtained using the python logistic regression classifier. Consistent results were obtained using the original Matlab implementation of the logistic regression classifier (see section S2 of the supplementary material). Single-subject results for each mask and trial estimate method are also reported in section S4 of the supplementary material.

**Figure 2.**
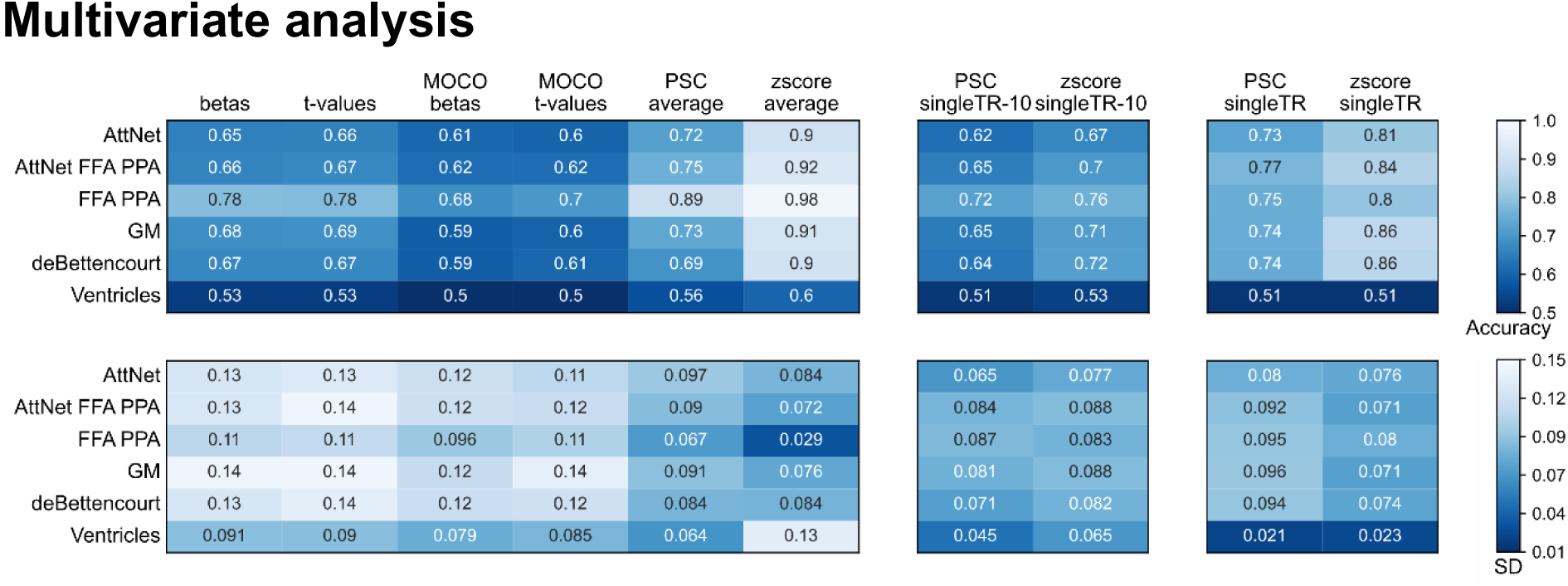
Python classification results. Upper table: mean accuracy scores across subjects for each mask and trial estimate method. Lower table: standard deviation of the accuracy scores across subjects. To facilitate comparisons, results were grouped according to the available number of samples. *Betas, t-values, PSC-avg, and zscore-avg* methods included 48 (or 40) samples, *PSC-singleTR-10* and *zscore-singleTR-10* resulted in 480 (or 400) samples, whereas *PSC-singleTR* and *zscore-singleTR* methods included 2400 (or 2000) samples per run.

The classification results of the ventricle masks showed average accuracy scores exceeding the conventional 0.5 chance level threshold (for a two-class problem) (Fig. 2-3). This effect was observed heterogeneously across subjects and trial estimate methods, and it was mainly due to motion-related artefacts in the BOLD time series and/or residual low-frequency drifts coupled with the task timing, which could not be removed from the data during the preprocessing steps (see Fig. 3 and Section 7 of the supplementary materials).

**Figure 3.**
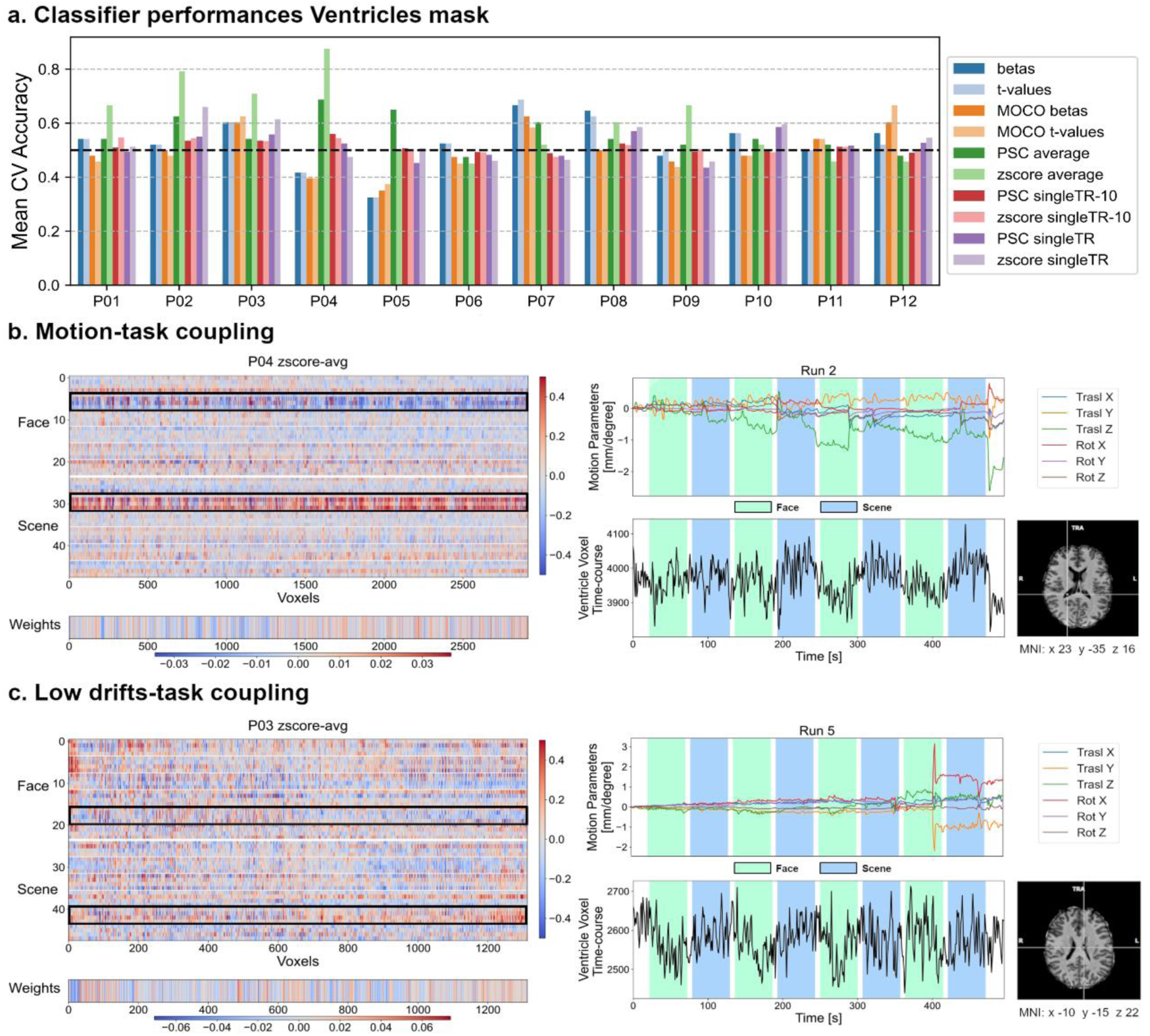
Pattern classification in the ventricles mask. a) Mean accuracy scores over cross-validation folds for each subject and trial estimate method. The black dashed line indicates the theoretical 0.5 chance level threshold. b) Example of motion-task coupling from run2 of participant P04. On the left, the patterns plot for the *zscore-avg* method, for which the mean classification accuracy was 0.875, and the corresponding classifier weights. Each row of the plot corresponds to a single trial. Trial estimates for Face and Scene are separated by the thick white middle line, and each subgroup of 4 rows corresponds to within run samples. On the right, representative time course from a ventricle voxel with motion-related artefacts (mainly due to Z translation). c) Example of low frequency drifts-task coupling from run 5 of participant P03. On the left, the patterns plot for the *zscore-avg* method, for which the mean classification accuracy was 0.71, and the corresponding classifier weights. Each row of the plot corresponds to a single trial. Trial estimates for Face and Scene are separated by the thick white middle line, and each subgroup of 4 rows corresponds to within run samples. On the right, a representative time course from a ventricle voxel with residual drifts coupled with the task timing (no motion contamination was present in this case).

Among the methods with a lower number of samples, the *zscore-avg* method exhibited the highest sensitivity to coupled noise drifts, reaching the average accuracy of 0.6 within the ventricle mask (Fig. 2, upper panel). This method also showed the highest standard deviation across subjects, consistent with the observation that coupling of low-frequency fluctuations with task timings was not present in all participants (Fig. 3a). Additionally, even though the data inspection unveiled, in some cases, the clear presence of a noise structure that should result in above chance classification, the *single-TR* methods did not strongly detect the noise bias. However, it was more visible when we included only the last 10 TRs of each task block, in which the BOLD signal should ideally have reached the plateau level. The inclusion of motion regressors in the GLM-based methods resulted in slightly lower accuracies, though the difference from the respective approaches without motion confounds was not statistically significant (as determined by a one-tailed paired Wilcoxon signed-rank test).

Regarding the classification performance across the different masks, the *FFA PPA mask* outperformed the other masking methods in terms of classification accuracy for all the trial estimate methods, except for the *single-TR* methods, reaching the average accuracy of 0.98 for the *zscore-avg* method, which also showed the lowest variance across subjects. Conversely, the *AttNet* mask, which did not include voxels from the FFA and PPA, performed on average worse than the other masking methods (Fig. 2). Among the trial estimates methods, GLM-based approaches showed less consistent results across subjects (Fig. 2, lower panel) and also lower accuracy scores when compared with the other small sample methods. The inclusion of motion regressors produced lower average accuracies compared to GLM-based approaches with only linear trends. One-tailed paired Wilcoxon signed-rank tests showed significantly lower accuracies for the *MOCO betas* method compared to *betas* in the *FFA PPA* mask and for the MOCO *t-values* method compared to *t-values* in both the *FFA PPA* and *GM* mask in the (p < 0.004, Bonferroni corrected n=12). Furthermore, in all method subgroups, the z-scoring normalisation approach resulted in the highest classification performances (Fig. 2).

### Simulation results

In the simulations with the fixed-order design, an average classification accuracy significantly higher than 0.5 was observed for the small sample methods when attempting to discriminate between the two classes in the null condition (i.e., no amplitude difference between the two conditions was imposed). This effect increased with an increase in the CNR value (Fig. 4). Furthermore, in contrast to the *t-values* and *betas* methods, the *PSC-avg* and *zscore-avg* methods showed a significant mean deviation of the accuracy distributions from the nominal chance level also for low CNR values (see section S6 of the supplementary materials for a detailed report of the statistical analysis). No significant deviation was found in the simulations with a random-order design for all trial estimation methods. The *singleTR* and *singleTR-10* methods did not show any significant change in the average accuracy with CNR or design type. Conversely, both *singleTR* and *singleTR-10* methods exhibited a significant increase in the variance of the accuracy scores distribution across CNR after Bonferroni correction (n=16; Levene’s test for fixed-order design: *PSC-singleTR-10,* F(3,3396)=72, p<0.003; *zscore-singleTR-10* F(3,3396)=158, p<0.003; *PSC-singleTR*, F(3,3396)=212, p<0.003; *zscore-singleTR*, F(3,3396)=295, p<0.003; Levene’s test for random-order design: *PSC-singleTR-10,* F(3,3396)=58, p<0.003; *zscore-singleTR-10* F(3,3396)=147, p<0.003; *PSC-singleTR*, F(3,3396)=199, p<0.003; *zscore-singleTR*, F(3,3396)=288, p<0.003). In contrast to the *singleTR* and *singleTR-10* methods, the *betas* and *t-values* methods showed a statistically significant reduction in variance at higher CNR, but only for the fixed-order design (Levene’s test for fixed-order design: *betas*, F(3,3396)=40, p<0.003, *t-values*, F(3,3396)=11, p<0.003). The accuracy distributions of *PSC-avg* and *zscore-avg* methods had approximately constant variance across CNR for both design types. Overall, the *single-TR* and *singleTR-10* methods showed the lowest variance in the accuracy score distributions as expected from the higher sample size (see section S6 of the supplementary materials).

**Figure 4.**
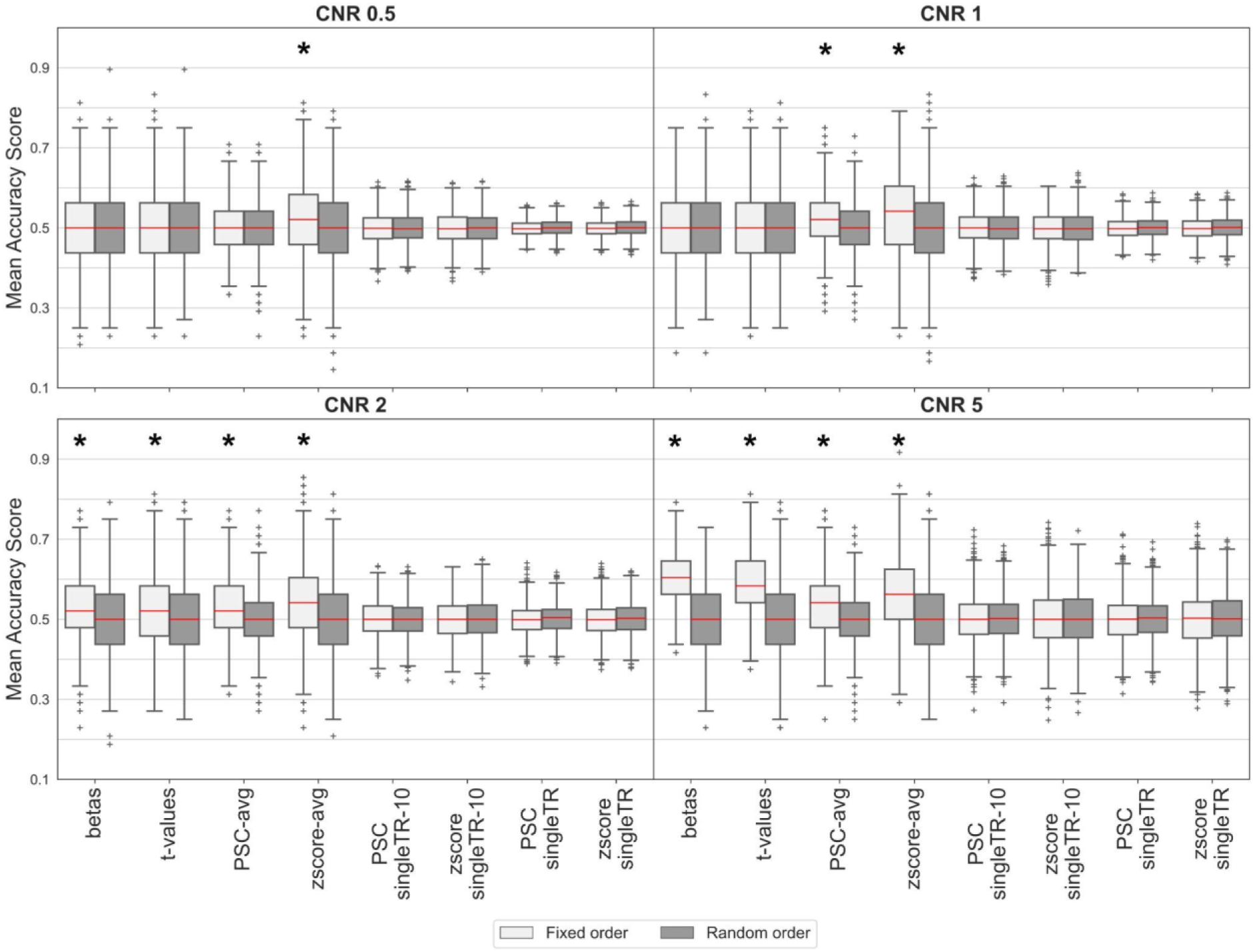
Simulation results. The box plots represent the accuracy score distributions obtained from 1000 simulations for each trial estimate method and CNR value. The mean accuracy obtained from the simulation with fixed-order design for the small sample methods increased with the CNR, whereas the *singleTR* methods showed only an increase in the variance of the accuracy scores distribution. Stars indicate accuracy distributions with a median significantly different than 0.5 (two-tailed one-sample Wilcoxon sign rank test, Bonferroni corrected n=64, see supplementary materials S6).

## Discussion

In this study, we investigated the influence of different trial estimation methods, brain masks and design choices on the classification accuracy of an LR classifier in a sustained attention paradigm. Participants were asked to focus on either the face or scene component in the composite (transparent) images that they were shown. An LR classifier was trained to distinguish between the two classes participants had to attend to (faces and scenes). We employed ventricular masks as control conditions for the classification task and performed simulations to assess the impact of the condition order on the classification performance. Generally, we showed that the small sample methods based on block averaging (*zscore-avg* and *PSC-avg*) exhibited the greatest sensitivity and that the FFA-PPA mask enabled the highest classification accuracy across the trial estimation methods. These results, however, have to be considered with caution, as we also showed that the analysis was influenced by multiple biases, mainly related to the study design. As no statistical significance of the accuracy scores could be assessed (as discussed below), accuracy scores were here only referred to as a relative measure of effect size across comparable conditions (Hebart & Baker, 2018).

The classification analysis conducted on the ventricle masks showed that, on average, the obtained accuracy scores exceeded the theoretical chance-level threshold of 0.5. Further examination of the data unveiled the presence of noise patterns intertwined with the experimental design. This noise-induced structure introduced a bias into the classification performance, resulting in the creation of artificial discriminative patterns within non-informative areas, such as the ventricles. As shown in section S7 of the supplementary material, these artefacts could potentially extend beyond the ventricular regions, impacting even areas that are genuinely associated with attention processes. Consequently, the differentiation between genuine effects and noise-induced patterns becomes almost impossible. In one of the reported example cases (Fig. 3b), the noise bias arose from task-related motion. Regressing out the motion parameters in this scenario would risk eliminating authentic effects or potentially retaining residual spurious signals that the classifier could still capture. Due to its high sensitivity to subtle class differences, motion-task coupling is even more problematic in MVPA than in mass univariate analysis and should be carefully investigated. Strict data exclusion criteria should also be considered. In a second instance, a low-frequency drift coupling was identified in the data (Fig. 2c), and this effect was also detrimental to the correct identification of real discriminative patterns. Indeed, the task frequency (∼0.008 Hz) was in a range commonly associated with low-frequency no-BOLD fluctuations (Gohel & Biswal, 2015). This poses challenges, as any attempt to temporally filter the data to eliminate these drifts could inadvertently eliminate task-related activity. Hence, it is imperative to carefully devise the experimental design to avoid analogous ambiguities during the data analysis process.

The present results undermine the correct interpretation of the absolute accuracy values obtained from classification analysis, as they might be inflated by structured noise. Nonetheless, relative comparisons remain useful to highlight properties of trial estimates and masking methods. Notably, among the small sample methods, the *zscore-avg* method (and secondly the *PSC-avg* method) exhibited the highest average accuracy scores and also the highest bias (Fig. 2). This could be explained considering that those methods incorporate within-run normalisation (i.e., z-scoring and PSC transformation) thus eliminating spurious shifts in the single-trial estimates across different runs (Lee & Kable, 2018). Indeed, a similar kind of within-run normalisation technique for trial estimates (e.g., mean centring) has been proven to significantly increase classification accuracy (Lee & Kable, 2018).

Single-trial estimation based on *betas*, *t-values* and *PSC-avg* methods encountered limitations in establishing an appropriate baseline level due to a limited number of time points available for baseline computation in the current design. For *betas* and *t-values* methods, the extended duration of the task block, combined with the short baseline period, might also have led to suboptimal model fitting, thereby yielding high variability of the trial estimates which, in turn, resulted in diminished accuracy scores. In fact, the BOLD signal of such a long task block might potentially present within block fluctuations attributable to e.g., changes in the attentional state of the participant and/or other physiological factors. Modelling this kind of signal as a single ideal response might be over-simplistic and hamper capturing the full variance of the BOLD signal. Consistently with a previous work by Stehr et al., the inclusion of motion confounds further lowered the classification accuracy (Stehr et al., 2023).

The *singleTR* methods, which resulted in the highest number of available samples, did not show the noise bias strongly while also potentially neglecting real effects. Previous studies (Stehr et al., 2023; Weaverdyck et al., 2020) discussed the implications of maximising the amount of data employed to train a classifier using single volumes instead of averaging approaches. Ultimately, these strategies increase the risk of poor classification performance if the signal-to-noise ratio (SNR) at the trial level is low, as might be the case in single-volume trials (Stehr et al., 2023). Furthermore, single trials within a block are highly correlated whereas learning algorithms assume that the input samples are independent and identically distributed (Huang et al., 2012).

Restricting the single trials to the last 10 TRs of each task block (*singleTR-10*) revealed an accuracy trend akin to the other small sample methods, despite the intrinsic lower SNR of the *singleTR* trials. This might be eventually due to the higher stability of the BOLD signal at the end of the task block.

The classification task was repeated using different masks to assess the influence of FFA and PPA spatial information. The *FFA PPA mask* showed the highest accuracy scores with a consistent performance trend across subjects and trial estimate methods. The only exceptions to this were the *singleTR* methods, which might be due to the lower sensitivity discussed above. Conversely, the *AttNet* mask showed on average the worst classification performances. In contrast with the findings of the previous work (DeBettencourt et al., 2015), this result suggests a significant contribution of the FFA and PPA spatial patterns to the classification accuracy. The previous work encompassed only a correlation analysis between the FFA-PPA mean activity difference and the classifier outputs, whereas our MVPA analysis could detect more nuanced variations in the spatial activity patterns that might remain concealed in a univariate analysis framework.

Furthermore, the inclusion of additional features using general brain masks (i.e., GM and deBettencourt masks) did not yield better performance. On the contrary, the introduction of many (and potentially non-informative) features might produce overfitting in the decoding analysis and lead to worse generalisation performances (Hawkins, 2004). A good choice of the features mask based on a priori anatomical information or functional localizer runs could help to have better discriminative power and also easier interpretability of the results (Weaverdyck et al., 2020, S. 20).

The analysis of the simulated data showed that the use of a fixed-order design could have introduced a bias in the classification performances evaluated on the real data. The presence of a fixed order in the presentation of the experimental conditions could indeed result in systematic effects in the data that the classifier might pick up. These results show that random order designs are crucial for MVPA analysis to prevent spurious classification success based on fixed-order effects.

In the simulations, averaging methods exhibited the highest sensitivity to the order effect as the average accuracies were significantly higher than 0.5 when no class effect was simulated in the data. For *betas* and *t-values* methods, the influence of the design on the classification accuracies was modulated by the CNR. This outcome is in line with previous results on real data where *betas* and *t-values* methods showed less sensitivity than averaging methods. We also observed an increase in the variance of the accuracy distributions with increasing CNR values in the *singleTR* and s*ingleTR-10* methods. This could not be attributed to a design influence as the effect was present for both the fixed- and the random-order simulated designs. However, this observation could be interpreted as an indication of the instability of these types of estimates, particularly when considering the results from the random-order design, which showed increasingly divergent accuracy scores even as data noise was reduced. Interestingly, *betas* and *t-values* methods showed a narrowing of the accuracy distribution with the CNR values only for the fixed-order design. This trend likely reflects a better fit of the GLM to the simulated data at higher CNR, capturing the order effect. Conversely, the *PSC-avg* and *zscore-avg* methods showed almost no difference in the variance of the accuracy distribution across CNR values, suggesting that these types of estimates exhibit more stable performance even in the presence of noisier data. Furthermore, in line with the results of the real data analyses, the *zscore-avg* method proved to be the most sensitive, detecting order effects at low CNR levels.

A direct implication of the simulation results was the impossibility of conducting any permutation or randomisation test to assess the significance of the obtained accuracy scores and perform hypothesis testing on the real data. Indeed, randomisation tests cannot be used when the design is fixed, and the presence of a short baseline period introduces dependencies across adjacent trial data making them not exchangeable for permutation procedures. In principle, within-run permutations would still be feasible considering only the last samples of each block (Valente et al., 2021) but the results would still be influenced by coupled noise in the analysed dataset. Permutation or randomisation tests are particularly needed when the final goal of the decoding analysis is the comparison of different classifier settings which might influence the shape of the distribution of cross-validated accuracy scores and make direct evaluation unfeasible. Indeed, the variance of cross-validated accuracy scores depends heavily on classifier type, cross-validation procedure, sample size and data dimensionality. All these factors influence the empirical threshold to be considered when evaluating statistical significance (Jamalabadi et al., 2016). In this work, we used relative changes in the accuracy scores to individuate potential biases into the decoding analysis and performed comparisons across similar decoding settings. However, the impossibility of performing any statistical assessment of the results and, primarily, the contamination of the data with different noise sources precluded a reliable evaluation of the actual effects on the analysed dataset. Nonetheless, our results provided several insights that could inform future advancements in the design of fMRI experiments for sustained attention training.

To optimise the classifier training in future applications of the paradigm, adapting the design by implementing a short block paradigm (shorter than the present 50 seconds) for the training runs should be considered. This would increase the number of training samples which is particularly problematic in real-time applications, while still enabling the use of robust averaging methods for more accurate trial estimates calculation. Furthermore, it would avoid the potential coupling of the task responses with low-frequency noise drifts observed in the current design. A longer baseline would also be advisable when using block designs to enable better response separation in adjacent blocks and also a more precise estimate of the baseline level. The order of the experimental conditions in the training runs should be randomised to avoid order bias in the classification results and allow statistical assessment. Based on our analysis, it could even be considered to restrict the classification only to FFA and PPA regions, as they showed the highest accuracies across participants.

## Conclusion

In this study, we examine how different trial estimation methods, brain masks, and design choices affect the classification accuracy of an LR classifier in a sustained attention paradigm. We identify the weaknesses and the limitations of the current design, which proved to be particularly prone to different biases when performing MVPA. We provide recommendations on how to improve the paradigm for future applications, pointing out the importance of careful study design to prevent biased results. Particularly, we strongly suggest a proper randomisation of the condition to avoid spurious inflation of the classification accuracy and to enable accurate statistical assessments through methods like permutation testing. Additionally, it is crucial to accurately define task timings, avoiding very low task repetition frequencies, to effectively differentiate task-related effects from noise fluctuations. Furthermore, we examine several properties of different trial estimation methods commonly used to conduct MVPA on fMRI data, showing the robustness of block averaging methods over single TR sampling. The selection of the most appropriate method should be guided by the specific requirements of the study design.

## Acknowledgements

We would like to thank Megan DeBettencourt and Nicholas Turk-Browne for the open exchange of information and scripts.

## References

Abdulrahman, H., & Henson, R. N. (2016). Effect of trial-to-trial variability on optimal event-related fMRI design: Implications for Beta-series correlation and multi-voxel pattern analysis. NeuroImage, 125, 756–766. 10.1016/j.neuroimage.2015.11.009

Abraham, A., Pedregosa, F., Eickenberg, M., Gervais, P., Mueller, A., Kossaifi, J., Gramfort, A., Thirion, B., & Varoquaux, G. (2014). Machine learning for neuroimaging with scikit-learn. Frontiers in Neuroinformatics, 8, 14. 10.3389/fninf.2014.00014

Al-Aidroos, N., Said, C. P., & Turk-Browne, N. B. (2012). Top-down attention switches coupling between low-level and high-level areas of human visual cortex. Proceedings of the National Academy of Sciences, 109(36), 14675– 14680. 10.1073/pnas.1202095109

Baldauf, D., & Desimone, R. (2014). Neural Mechanisms of Object-Based Attention. Science, 344(6182), 424–427. 10.1126/science.1247003

Baucom, L. B., Wedell, D. H., Wang, J., Blitzer, D. N., & Shinkareva, S. V. (2012). Decoding the neural representation of affective states. NeuroImage, 59(1), 718–727. 10.1016/j.neuroimage.2011.07.037

Bressler, R. A., Raible, S., Lührs, M., Tier, R., Goebel, R., & Linden, D. E. (2023). No threat: Emotion regulation neurofeedback for police special forces recruits. Neuropsychologia, 190, 108699. 10.1016/j.neuropsychologia.2023.108699

Chee, M. W. L., Tan, J. C., Parimal, S., & Zagorodnov, V. (2010). Sleep deprivation and its effects on object-selective attention. NeuroImage, 49(2), 1903–1910. 10.1016/j.neuroimage.2009.08.067

Chiu, Y.-C., Esterman, M., Han, Y., Rosen, H., & Yantis, S. (2011). Decoding Task-based Attentional Modulation during Face Categorization. Journal of Cognitive Neuroscience, 23(5), 1198–1204. 10.1162/jocn.2010.21503

Cohen, J. D., Daw, N., Engelhardt, B., Hasson, U., Li, K., Niv, Y., Norman, K. A., Pillow, J., Ramadge, P. J., Turk-Browne, N. B., & Willke, T. L. (2017). Computational approaches to fMRI analysis. Nature Neuroscience, 20(3), Article 3. 10.1038/nn.4499

Cohen, O., Koppel, M., Malach, R., & Friedman, D. (2014). Controlling an avatar by thought using real-time fMRI. Journal of Neural Engineering, 11(3), 035006. 10.1088/1741-2560/11/3/035006

Cox, D. D., & Savoy, R. L. (2003). Functional magnetic resonance imaging (fMRI) „brain reading“: Detecting and classifying distributed patterns of fMRI activity in human visual cortex. NeuroImage, 19(2), 261–270. 10.1016/S1053-8119(03)00049-1

Craddock, R. C., James, G. A., Holtzheimer, P. E., Hu, X. P., & Mayberg, H. S. (2012). A whole brain fMRI atlas generated via spatially constrained spectral clustering. Human Brain Mapping, 33(8), 1914–1928. 10.1002/hbm.21333

De Martino, F., Valente, G., Staeren, N., Ashburner, J., Goebel, R., & Formisano, E. (2008). Combining multivariate voxel selection and support vector machines for mapping and classification of fMRI spatial patterns. NeuroImage, 43(1), 44–58. 10.1016/j.neuroimage.2008.06.037

DeBettencourt, M. T., Cohen, J. D., Lee, R. F., Norman, K. A., & Turk-Browne, N. B. (2015). Closed-loop training of attention with real-time brain imaging. Nature Neuroscience, 18(3), 470–478. 10.1038/nn.3940

Downing, P., Liu, J., & Kanwisher, N. (2001). Testing cognitive models of visual attention with fMRI and MEG. Neuropsychologia, 39(12), 1329–1342. 10.1016/S0028-3932(01)00121-X

Formisano, E., De Martino, F., & Valente, G. (2008). Multivariate analysis of fMRI time series: Classification and regression of brain responses using machine learning. Magnetic Resonance Imaging, 26(7), 921–934. 10.1016/j.mri.2008.01.052

Fox, M. D., Corbetta, M., Snyder, A. Z., Vincent, J. L., & Raichle, M. E. (2006). Spontaneous neuronal activity distinguishes human dorsal and ventral attention systems. Proceedings of the National Academy of Sciences of the United States of America, 103(26), 10046–10051. 10.1073/pnas.0604187103

Fox, M. D., Snyder, A. Z., Vincent, J. L., Corbetta, M., Van Essen, D. C., & Raichle, M. E. (2005). The human brain is intrinsically organized into dynamic, anticorrelated functional networks. Proceedings of the National Academy of Sciences of the United States of America, 102(27), 9673–9678. 10.1073/pnas.0504136102

Friston, K. J., Fletcher, P., Josephs, O., Holmes, A., Rugg, M. D., & Turner, R. (1998). Event-Related fMRI: Characterizing Differential Responses. NeuroImage, 7(1), 30–40. 10.1006/nimg.1997.0306

Gohel, S. R., & Biswal, B. B. (2015). Functional Integration Between Brain Regions at Rest Occurs in Multiple-Frequency Bands. Brain Connectivity, 5(1), 23–34. 10.1089/brain.2013.0210

Guo, Y., Liu, Y., Oerlemans, A., Lao, S., Wu, S., & Lew, M. S. (2016). Deep learning for visual understanding: A review. Neurocomputing, 187, 27–48. 10.1016/j.neucom.2015.09.116

Hawkins, D. M. (2004). The Problem of Overfitting. 1–12.

Haynes, J.-D. (2015). A Primer on Pattern-Based Approaches to fMRI: Principles, Pitfalls, and Perspectives. Neuron, 87(2), 257–270. 10.1016/j.neuron.2015.05.025

Hebart, M. N., & Baker, C. I. (2018). Deconstructing multivariate decoding for the study of brain function. NeuroImage, 180, 4–18. 10.1016/j.neuroimage.2017.08.005

Huang, X., Jin, G., & Ruan, W. (2012). Machine Learning Basics. In X. Huang, G. Jin, & W. Ruan (Hrsg.), Machine Learning Safety (S. 3–13). Springer Nature. 10.1007/978-981-19-6814-3_1

Jamalabadi, H., Alizadeh, S., Schönauer, M., Leibold, C., & Gais, S. (2016). Classification based hypothesis testing in neuroscience: Below-chance level classification rates and overlooked statistical properties of linear parametric classifiers. Human Brain Mapping, 37(5), 1842–1855. 10.1002/hbm.23140

LaConte, S. M., Peltier, S. J., & Hu, X. P. (2007). Real-time fMRI using brain-state classification. Human Brain Mapping, 28(10), 1033–1044. 10.1002/hbm.20326

Langner, R., & Eickhoff, S. B. (2013). Sustaining attention to simple tasks: A meta-analytic review of the neural mechanisms of vigilant attention. Psychological Bulletin, 139(4), 870–900. 10.1037/a0030694

Lecun, Y., Bengio, Y., & Hinton, G. (2015). Deep learning. Nature Reviews Neuroscience, 521(7553), 436–444. 10.1038/nature14539

Lee, S., & Kable, J. W. (2018). Simple but robust improvement in multivoxel pattern classification. PLOS ONE, 13(11), e0207083. 10.1371/journal.pone.0207083

Levene, H. (1960). Contributions to probability and statistics. Essays in honor of Harold Hotelling, 278, 292.

Liang, Y., Bo, K., Meyyappan, S., & Ding, M. (2023). Decoding fMRI Data: A Comparison Between Support Vector Machines and Deep Neural Networks (S. 2023.05.30.542882). bioRxiv. 10.1101/2023.05.30.542882

Madsen, K. H., Churchill, N. W., & Mørup, M. (2017). Quantifying functional connectivity in multi-subject fMRI data using component models. Human Brain Mapping, 38(2), 882–899. 10.1002/hbm.23425

Madsen, K. H., Krohne, L. G., Cai, X. L., Wang, Y., & Chan, R. C. K. (2018). Perspectives on Machine Learning for Classification of Schizotypy Using fMRI Data. Schizophrenia Bulletin, 44(2), S480–S490. 10.1093/schbul/sby026

Mahmoudi, A., Takerkart, S., Regragui, F., Boussaoud, D., & Brovelli, A. (2012). Multivoxel pattern analysis for FMRI data: A review. Computational and Mathematical Methods in Medicine, 2012, 961257. 10.1155/2012/961257

Mandelkow, H., de Zwart, J. A., & Duyn, J. H. (2016). Linear Discriminant Analysis Achieves High Classification Accuracy for the BOLD fMRI Response to Naturalistic Movie Stimuli. Frontiers in Human Neuroscience, 10. https://www.frontiersin.org/articles/10.3389/fnhum.2016.00128

McDonald, A. R., Muraskin, J., Dam, N. T. V., Froehlich, C., Puccio, B., Pellman, J., Bauer, C. C. C., Akeyson, A., Breland, M. M., Calhoun, V. D., Carter, S., Chang, T. P., Gessner, C., Gianonne, A., Giavasis, S., Glass, J., Homann, S., King, M., Kramer, M., … Craddock, R. C. (2017). The real-time fMRI neurofeedback based stratification of Default Network Regulation Neuroimaging data repository. NeuroImage, 146, 157–170. 10.1016/j.neuroimage.2016.10.048

Misaki, M., Kim, Y., Bandettini, P. A., & Kriegeskorte, N. (2010). Comparison of multivariate classifiers and response normalizations for pattern-information fMRI. NeuroImage, 53(1), 103–118. 10.1016/j.neuroimage.2010.05.051

Mumford, J. A., Davis, T., & Poldrack, R. A. (2014). The impact of study design on pattern estimation for single-trial multivariate pattern analysis. NeuroImage, 103, 130–138. 10.1016/j.neuroimage.2014.09.026

Mur, M., Bandettini, P. A., & Kriegeskorte, N. (2009). Revealing representational content with pattern-information fMRI - An introductory guide. Social Cognitive and Affective Neuroscience, 4(1), 101–109. 10.1093/scan/nsn044

O’Craven, K. M., Downing, P. E., & Kanwisher, N. (1999). fMRI evidence for objects as the units of attentional selection. Nature, 401(6753), 584–587. 10.1038/44134

Pedregosa, F., Varoquaux, G., Gramfort, A., Michel, V., Thirion, B., Grisel, O., Blondel, M., Prettenhofer, P., Weiss, R., Dubourg, V., Vanderplas, J., Passos, A., Cournapeau, D., Brucher, M., Perrot, M., & Duchesnay, É. (2011). Scikit-learn: Machine Learning in Python. Journal of Machine Learning Research, 12(85), 2825–2830.

Posner, M. I., & Dehaene, S. (1994). Attentional networks. Trends in Neurosciences, 17(2), 75–79. 10.1016/0166-2236(94)90078-7

Power, J. D., Cohen, A. L., Nelson, S. M., Wig, G. S., Barnes, K. A., Church, J. A., Vogel, A. C., Laumann, T. O., Miezin, F. M., Schlaggar, B. L., & Petersen, S. E. (2011). Functional Network Organization of the Human Brain. Neuron, 72(4), 665–678. 10.1016/j.neuron.2011.09.006

Prince, J. S., Charest, I., Kurzawski, J. W., Pyles, J. A., Tarr, M. J., & Kay, K. N. (2022). Improving the accuracy of single-trial fMRI response estimates using GLMsingle. eLife, 11, e77599. 10.7554/eLife.77599

Reddy, L., Moradi, F., & Koch, C. (2007). Top–down biases win against focal attention in the fusiform face area. NeuroImage, 38(4), 730–739. 10.1016/j.neuroimage.2007.08.006

Sitaram, R., Lee, S., Ruiz, S., Rana, M., Veit, R., & Birbaumer, N. (2011). Real-time support vector classification and feedback of multiple emotional brain states. NeuroImage, 56(2), 753–765. 10.1016/j.neuroimage.2010.08.007

Smith, S. M., Miller, K. L., Salimi-Khorshidi, G., Webster, M., Beckmann, C. F., Nichols, T. E., Ramsey, J. D., & Woolrich, M. W. (2011). Network modelling methods for FMRI. NeuroImage, 54(2), 875–891. 10.1016/j.neuroimage.2010.08.063

Smucny, J., Shi, G., & Davidson, I. (2022). Deep Learning in Neuroimaging: Overcoming Challenges With Emerging Approaches. Frontiers in Psychiatry, 13. https://www.frontiersin.org/articles/10.3389/fpsyt.2022.912600

Sorger, B., & Goebel, R. (2020). Real-time fMRI for brain-computer interfacing. In Handbook of Clinical Neurology (Bd. 168, S. 289–302). Elsevier. 10.1016/B978-0-444-63934-9.00021-4

Stehr, D. A., Garcia, J. O., Pyles, J. A., & Grossman, E. D. (2023). Optimizing multivariate pattern classification in rapid event-related designs. Journal of Neuroscience Methods, 387, 109808. 10.1016/j.jneumeth.2023.109808

Taschereau-Dumouchel, V., Cortese, A., Chiba, T., Knotts, J. D., Kawato, M., & Lau, H. (2018). Towards an unconscious neural reinforcement intervention for common fears. Proceedings of the National Academy of Sciences, 115(13), 3470–3475. 10.1073/pnas.1721572115

Valente, G., Castellanos, A. L., Hausfeld, L., De Martino, F., & Formisano, E. (2021). Cross-validation and permutations in MVPA: Validity of permutation strategies and power of cross-validation schemes. NeuroImage, 238, 118145. 10.1016/j.neuroimage.2021.118145

Wang, L., Lei, Y., Zeng, Y., Tong, L., & Yan, B. (2013). Principal Feature Analysis: A Multivariate Feature Selection Method for fMRI Data. Computational and Mathematical Methods in Medicine, 2013, e645921. 10.1155/2013/645921

Wang, X., Liang, X., Jiang, Z., Nguchu, B. A., Zhou, Y., Wang, Y., Wang, H., Li, Y., Zhu, Y., Wu, F., Gao, J., & Qiu, B. (2020). Decoding and mapping task states of the human brain via deep learning. Human Brain Mapping, 41(6), 1505– 1519. 10.1002/hbm.24891

Weaverdyck, M. E., Lieberman, M. D., & Parkinson, C. (2020). Tools of the Trade Multivoxel pattern analysis in fMRI: A practical introduction for social and affective neuroscientists. Social Cognitive and Affective Neuroscience, 15(4), 487–509. 10.1093/scan/nsaa057

Welvaert, M., & Rosseel, Y. (2013). On the Definition of Signal-To-Noise Ratio and Contrast-To-Noise Ratio for fMRI Data. PLOS ONE, 8(11), e77089. 10.1371/journal.pone.0077089

Wojciulik, E., Kanwisher, N., & Driver, J. (1998). Covert Visual Attention Modulates Face-Specific Activity in the Human Fusiform Gyrus: fMRI Study. Journal of Neurophysiology, 79(3), 1574–1578. 10.1152/jn.1998.79.3.1574

Yi, D.-J., Kelley, T. A., Marois, R., & Chun, M. M. (2006). Attentional modulation of repetition attenuation is anatomically dissociable for scenes and faces. Brain Research, 1080(1), 53–62. 10.1016/j.brainres.2006.01.090

