## supplementary for "Navigating the Maze: Identifying Potential Pitfalls in Attention State Classification from fMRI Brain Patterns"

### Supplementary Materials

#### S1. Masks details

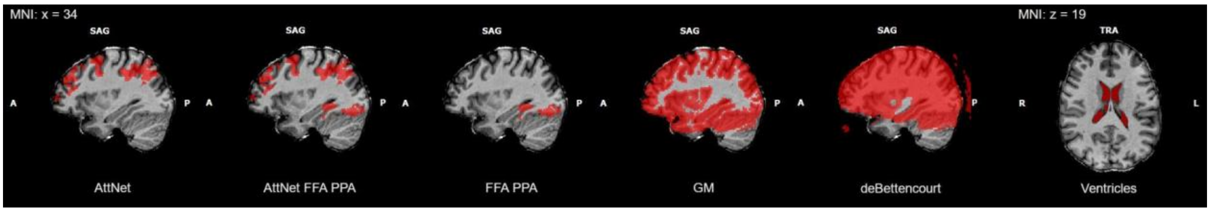


The figure shows example masks from participant P01 in MNI space. The *AttNet* mask, obtained based on the attention network localizer run, mainly included parietal and frontal regions involved in the attention task, such as anterior midcingulate cortex (aMCC), the frontal eye field (FEF), the inferior frontal junction (IFJ) and the intraparietal sulcus (IPS). The *FFA PPA* mask was created from the probabilistic functional atlas presented in (Rosenke et al., 2021) selecting the following sub-regions: *lh_CoS_places, lh_IOG_faces, lh_mFus_faces, lh_pFus_faces, rh_CoS_places, rh_IOG_faces, rh_mFus_faces, rh_pFus_faces*. The *AttNet FFA PPA* mask was obtained by combining the *AttNet* mask and the *FFA PPA* mask. The *GM* mask consisted of all cortical, subcortical and cerebellar grey matter included in the functional field of view. The *deBettencourt* mask, obtained following the procedure of the previous study (DeBettencourt et al., 2015), covered most of the brain (grey matter, white matter and CSF) with some asymmetries in the subcortical and temporal regions, which were sometimes partially excluded. Most notably, the skull and eyes were often included. *Ventricles* masks were created from individual structural images, selecting connected areas within the ventricles with intensity values lower than 40.
Since the masks were generated individually for each subject (with the only exception of the *FFA PPA* mask), a variable number of voxels was employed in the MVPA across subjects for the same mask type. The table below shows the dimensions information for each mask type and average FFA PPA coverage:

| | AttNet | AttNet FFA PPA | FFA PPA | GM | deBettencourt | Ventricles |
|---|---|---|---|---|---|---|
| mean (# voxels) | 46817 | 49466 | 2649 | 169224 | 193040 | 2186 |
| min (# voxels) | 16910 | 19559 | 2649 | 161213 | 179580 | 473 |
| max (# voxels) | 72857 | 75506 | 2649 | 184149 | 207868 | 5472 |
| FFA PPA average coverage | 0% | 100% | 100% | 88% | 92% | 0% |

#### S2. Classification performances logistic regression (Matlab)

Accuracy

| | betas | t-values | MOCO betas | MOCO t-values | PSC average | zscore average | PSC singleTR-10 | zscore singleTR-10 | PSC singleTR | zscore singleTR |
|---|---|---|---|---|---|---|---|---|---|---|
| AttNet | 0.63 | 0.65 | 0.58 | 0.58 | 0.72 | 0.87 | 0.61 | 0.64 | 0.73 | 0.78 |
| AttNet FFA PPA | 0.65 | 0.67 | 0.6 | 0.58 | 0.73 | 0.89 | 0.63 | 0.67 | 0.77 | 0.82 |
| FFA PPA | 0.78 | 0.78 | 0.7 | 0.71 | 0.89 | 0.97 | 0.71 | 0.75 | 0.77 | 0.82 |
| GM | 0.68 | 0.68 | 0.57 | 0.57 | 0.7 | 0.89 | 0.62 | 0.7 | 0.76 | 0.84 |
| deBettencourt | 0.66 | 0.62 | 0.57 | 0.56 | 0.7 | 0.89 | 0.62 | 0.69 | 0.75 | 0.83 |
| Ventricles | 0.52 | 0.51 | 0.49 | 0.48 | 0.56 | 0.61 | 0.52 | 0.53 | 0.51 | 0.51 |

SD

| | betas | t-values | MOCO betas | MOCO t-values | PSC average | zscore average | PSC singleTR-10 | zscore singleTR-10 | PSC singleTR | zscore singleTR |
|---|---|---|---|---|---|---|---|---|---|---|
| AttNet | 0.13 | 0.12 | 0.095 | 0.14 | 0.1 | 0.086 | 0.06 | 0.071 | 0.08 | 0.062 |
| AttNet FFA PPA | 0.14 | 0.14 | 0.11 | 0.13 | 0.081 | 0.097 | 0.078 | 0.084 | 0.085 | 0.076 |
| FFA PPA | 0.11 | 0.09 | 0.098 | 0.1 | 0.075 | 0.034 | 0.089 | 0.08 | 0.072 | 0.072 |
| GM | 0.12 | 0.11 | 0.11 | 0.11 | 0.093 | 0.095 | 0.075 | 0.083 | 0.097 | 0.072 |
| deBettencourt | 0.11 | 0.12 | 0.12 | 0.11 | 0.071 | 0.096 | 0.072 | 0.076 | 0.091 | 0.074 |
| Ventricles | 0.081 | 0.083 | 0.095 | 0.085 | 0.056 | 0.13 | 0.045 | 0.063 | 0.023 | 0.025 |

The table reports the average accuracy scores across all subjects obtained from the cross-validation procedure using the Matlab implementation of the logistic regression classifier (DeBettencourt et al., 2015). On average slightly lower accuracy scores but the same trends of the python *sklearn* logistic regression were observed.

#### S3. Physiological data quality

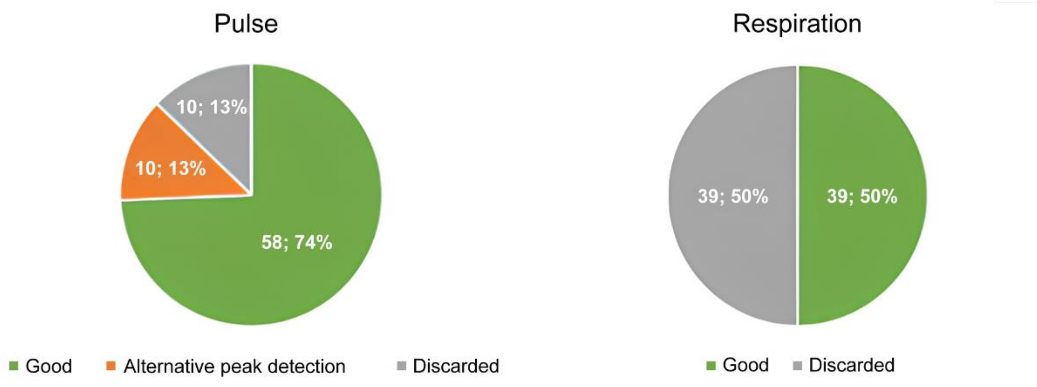


The figure above shows a summary of the physiological data quality. Data of the training runs for all participants and sessions included a total of 78 runs. Data of good quality could be used without further adaptations of the peak extraction methods. For the cardiac data, in some cases, data quality could be improved by using an alternative peak detection method. Data of insufficient quality for analysis had to be discarded.

#### S4. Single subject classification results (python)

The following plots show single-subject average accuracy over cross-validation folds for each mask and trial estimate method, using the python *sklearn* logistic regression classifier.

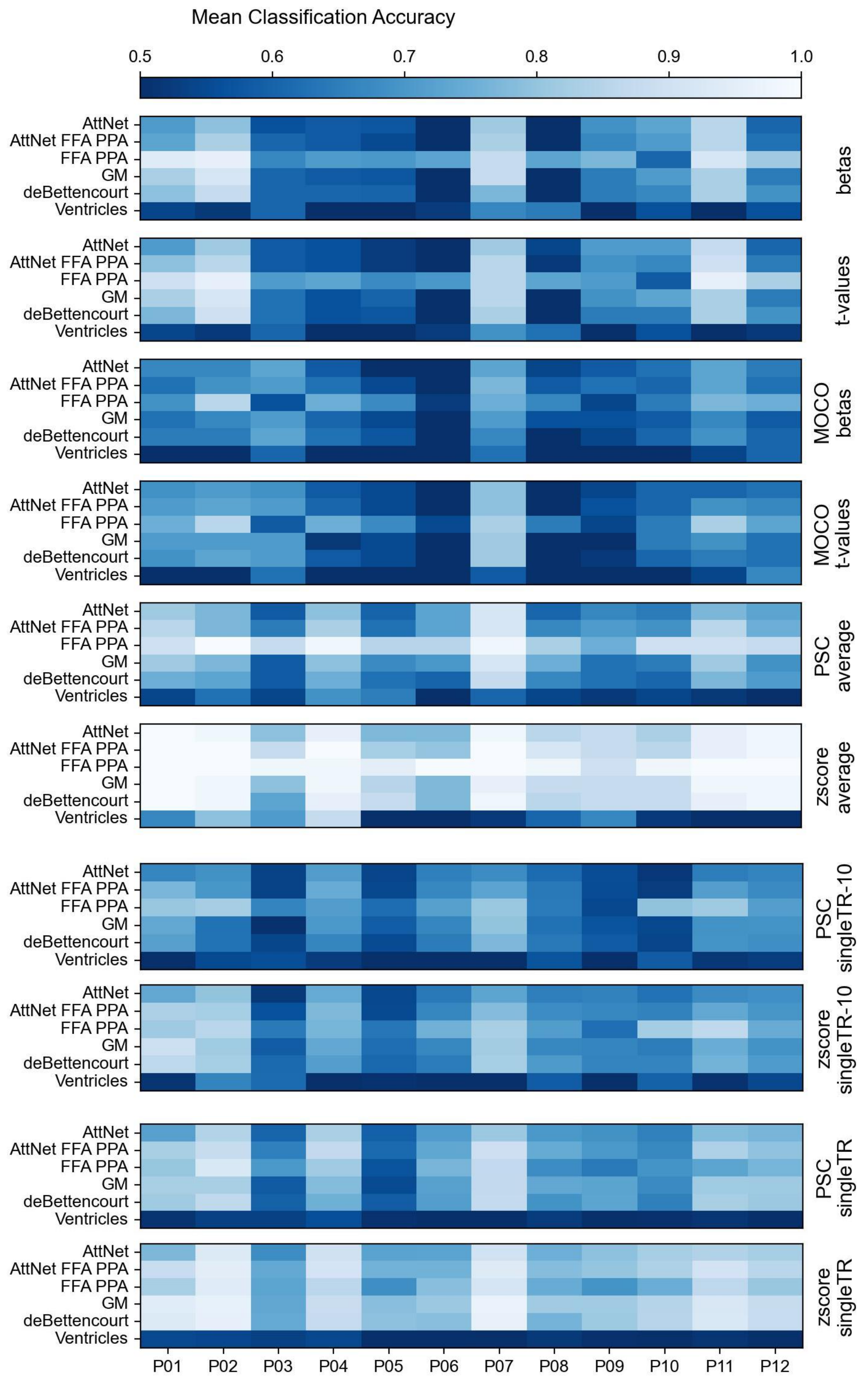

Mean Classification Accuracy
0.5
0.6
0.7
0.8
0.9
1.0
AttNet
AttNet FFA PPA
FFA PPA
GM
deBettencourt
Ventricles
betas
t-values
MOCO betas
MOCO t-values
PSC average
zscore average
PSC singleTR-10
zscore singleTR-10
PSC singleTR
zscore singleTR
P01
P02
P03
P04
P05
P06
P07
P08
P09
P10
P11
P12

#### S5 Simulations example

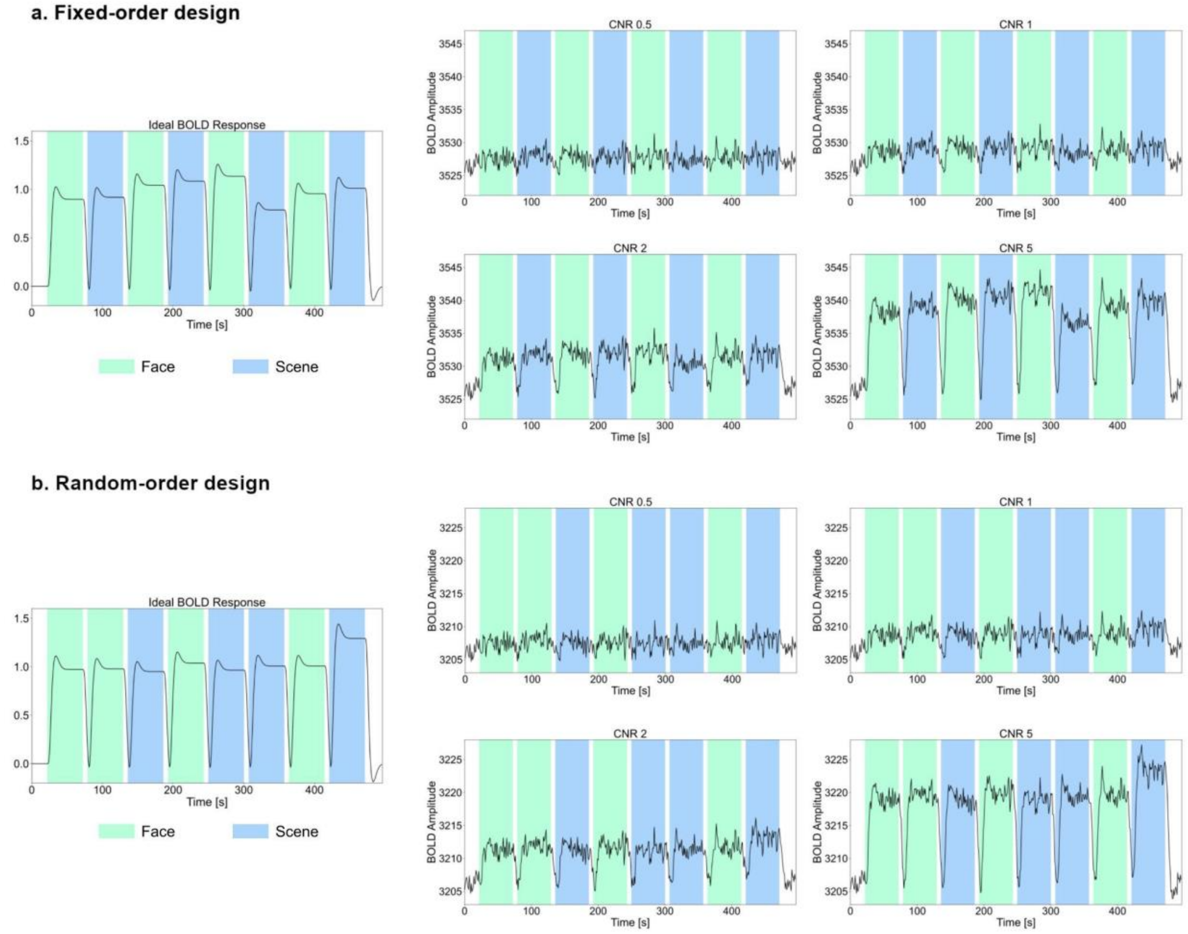


Simulated time-course examples at different CNR. a) Time-course example with fixed-order design as in the real data. On the left, the ideal time course with random amplitude modulation. On the right, the final time-course obtained by combining the ideal time-course and the autocorrelated noise with CNR scaling (Welvaert & Rosseel, 2013). b) Time-course example with random-order design. On the left, the ideal time course with random amplitude modulation. On the right, the final time-course obtained combining the ideal time-course and the autocorrelated noise with CNR scaling. The number of blocks per condition was kept constant in each simulated run, whereas the order of the conditions was changed at random.

#### S6. Simulations: statistical analysis

| | betas | | | t-values | | | PSC-avg | | | zscore-avg | | |
|---|---|---|---|---|---|---|---|---|---|---|---|---|
| **FIXED** | median | z | p | median | z | p | median | z | p | median | z | p |
| CNR 0.5 | 0.50 | -0.83 | 0.40 | 0.50 | -0.60 | 0.55 | 0.50 | 3.21 | 0.001 | 0.52 | 4.55 | <0.001 |
| CNR 1 | 0.50 | -1.32 | 0.19 | 0.50 | 0.06 | 0.95 | 0.52 | 8.53 | <0.001 | 0.54 | 10.29 | <0.001 |
| CNR 2 | 0.52 | 10.29 | <0.001 | 0.52 | 6.49 | <0.001 | 0.52 | 12.01 | <0.001 | 0.54 | 13.80 | <0.001 |
| CNR 5 | 0.60 | 26.57 | <0.001 | 0.58 | 24.55 | <0.001 | 0.54 | 13.82 | <0.001 | 0.56 | 15.52 | <0.001 |
| **RANDOM** | | | | | | | | | | | | |
| CNR 0.5 | 0.50 | 0.03 | 0.98 | 0.50 | 0.92 | 0.36 | 0.50 | 0.88 | 0.38 | 0.50 | 0.69 | 0.49 |
| CNR 1 | 0.50 | -0.20 | 0.84 | 0.50 | 0.97 | 0.33 | 0.50 | 0.28 | 0.78 | 0.50 | 0.78 | 0.44 |
| CNR 2 | 0.50 | 1.27 | 0.20 | 0.50 | 1.87 | 0.06 | 0.50 | 0.48 | 0.63 | 0.50 | 1.48 | 0.14 |
| CNR 5 | 0.50 | -0.22 | 0.83 | 0.50 | -0.41 | 0.68 | 0.50 | 1.07 | 0.28 | 0.50 | 1.41 | 0.16 |

| | PSC-singleTR-10 | | | zscore-singleTR-10 | | | PSC-singleTR | | | zscore-singleTR | | |
|---|---|---|---|---|---|---|---|---|---|---|---|---|
| **FIXED** | median | z | p | median | z | p | median | z | p | median | z | p |
| CNR 0.5 | 0.50 | -0.45 | 0.65 | 0.50 | -1.04 | 0.30 | 0.50 | -2.42 | 0.02 | 0.50 | -1.97 | 0.05 |
| CNR 1 | 0.50 | 0.10 | 0.92 | 0.50 | -0.47 | 0.64 | 0.50 | -1.83 | 0.07 | 0.50 | -1.81 | 0.07 |
| CNR 2 | 0.50 | -0.08 | 0.94 | 0.50 | -0.38 | 0.70 | 0.50 | -0.64 | 0.52 | 0.50 | -1.02 | 0.31 |
| CNR 5 | 0.50 | -0.04 | 0.97 | 0.50 | 0.20 | 0.84 | 0.50 | -0.26 | 0.79 | 0.50 | -0.45 | 0.65 |
| **RANDOM** | | | | | | | | | | | | |
| CNR 0.5 | 0.50 | -0.76 | 0.45 | 0.50 | -0.77 | 0.44 | 0.50 | 0.70 | 0.49 | 0.50 | 1.07 | 0.28 |
| CNR 1 | 0.50 | -0.23 | 0.81 | 0.50 | -0.64 | 0.52 | 0.50 | 1.01 | 0.31 | 0.50 | 1.26 | 0.21 |
| CNR 2 | 0.50 | 0.45 | 0.65 | 0.50 | 0.41 | 0.68 | 0.50 | 1.38 | 0.17 | 0.50 | 1.24 | 0.22 |
| CNR 5 | 0.50 | 0.38 | 0.71 | 0.50 | 0.90 | 0.37 | 0.50 | 0.86 | 0.39 | 0.50 | 1.03 | 0.30 |

The table shows the results of the two-tailed Wolcoxon sign rank performed on the accuracy values obtained from the 1000 simulations for each trial estimate method and CNR to test that the median of the distributions was different from the theoretical chance level of 0.5. Coloured cells indicate significant results after Bonferroni correction (n=64). Results for all small sample size methods were significant for CNR 2 and 5, *betas* exhibited the highest effect with a median of 0.6. PSC-avg and zscore-avg showed significant results also at CNR 0.5 and 1.

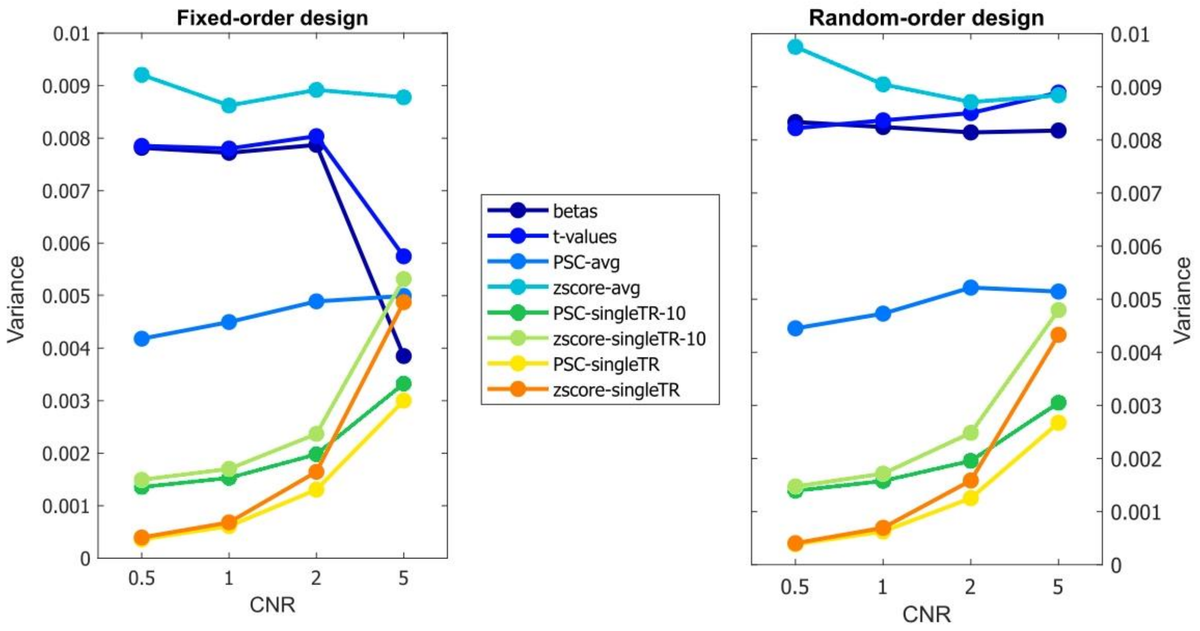


The *singleTR* methods did not show any change in average accuracy with CNR or design type. On the other hand, both *singleTR* methods exhibited a significant increase in the variance of the accuracy scores distribution across CNR after Bonferroni correction (n=16; Levene's test for fixed-order design: *PSC-singleTR*, F(3,3396)=212, p<0.003, *zscore-singleTR*, F(3,3396)=295, p<0.003; Levene's test for random-order design: *PSC-singleTR* , F(3,3396)=198, p<0.003, *zscore-singleTR*, F(3,3396)=288, p<0.003). Interestingly, both betas and t-values methods showed an opposite trend only for the fixed-order design (Levene's test for fixed-order design: *betas*, F(3,3396)=40, p<0.003, *tvals*, F(3,3396)=11, p<0.003). The accuracy distributions of *PSC-avg* and *zscore-avg* methods had approximately constant variance across CNR for both design types.

#### S7. Noise-task coupling example

The figure below shows the seed-based correlation map from run 2 of participant P04. The seed time-course was chosen within the right ventricle (MNI coordinates: x=23, y=-35, z=16). As reported in the main manuscript (Fig. 2b), motion-task coupling was observed within the ventricles, where no task-related activity was expected. The correlation map (p<0.001) proved that the coupling pattern extended to the parietal regions involved in the attention process, contaminating the actual task effects.

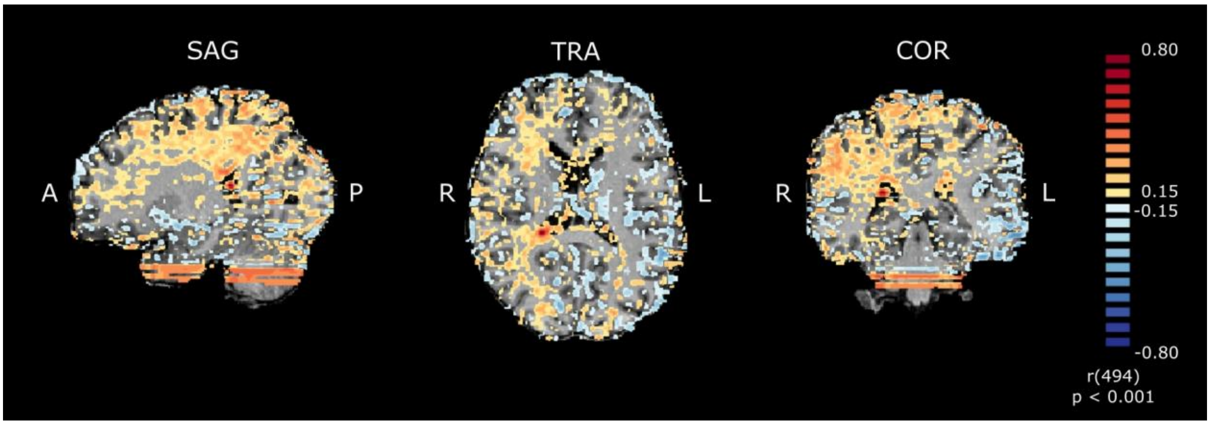


### References


DeBettencourt, M. T., Cohen, J. D., Lee, R. F., Norman, K. A., & Turk-Browne, N. B. (2015). Closed-loop training of attention with real-time brain imaging. *Nature Neuroscience*, *18*(3), 470–478. https://doi.org/10.1038/nn.3940

Rosenke, M., van Hoof, R., van den Hurk, J., Grill-Spector, K., & Goebel, R. (2021). A Probabilistic Functional Atlas of Human Occipito-Temporal Visual Cortex. *Cerebral Cortex*, *31*(1), 603–619. https://doi.org/10.1093/cercor/bhaa246

Welvaert, M., & Rosseel, Y. (2013). On the Definition of Signal-To-Noise Ratio and Contrast-To-Noise Ratio for fMRI Data. *PLOS ONE*, *8*(11), e77089. https://doi.org/10.1371/journal.pone.0077089
